# Biomaterial-based 3D human lung models replicate pathological characteristics of early pulmonary fibrosis

**DOI:** 10.1101/2025.02.12.637970

**Authors:** Alicia E. Tanneberger, Rachel Blomberg, Anton D. Kary, Andrew Lu, David W.H. Riches, Chelsea M. Magin

## Abstract

Idiopathic pulmonary fibrosis (IPF) is a progressive and incurable lung disease characterized by tissue scarring that disrupts gas exchange. Epithelial cell dysfunction, fibroblast activation, and excessive extracellular matrix deposition drive this pathology that ultimately leads to respiratory failure. Mechanistic studies have shown that repeated injury to alveolar epithelial cells initiates an aberrant wound-healing response in surrounding fibroblasts through secretion of mediators like transforming growth factor-β, yet the precise biological pathways contributing to disease progression are not fully understood. To better study these interactions there is a critical need for lung models that replicate the cellular heterogeneity, geometry, and biomechanics of the distal lung microenvironment. In this study, induced pluripotent stem cell-derived alveolar epithelial type II (iATII) cells and human pulmonary fibroblasts were arranged to replicate human lung micro-architecture and embedded in soft or stiff poly(ethylene glycol) norbornene (PEG-NB) hydrogels that recapitulated the mechanical properties of healthy and fibrotic lung tissue, respectively. The co-cultured cells were then exposed to pro-fibrotic biochemical cues, including inflammatory cytokines and growth factors. iATIIs and fibroblasts exhibited differentiation pathways and gene expression patterns consistent with trends observed during IPF progression *in vivo*. A design of experiments statistical analysis identified stiff hydrogels combined with pro-fibrotic biochemical cue exposure as the most effective condition for modeling fibrosis *in vitro*. Finally, treatment with Nintedanib, one of only two Food and Drug Administration (FDA)-approved drugs for IPF, was assessed. Treatment reduced fibroblast activation, as indicated by downregulation of key activation genes, and upregulated several epithelial genes. These findings demonstrate that human 3D co-culture models hold tremendous potential for advancing our understanding of IPF and identifying novel therapeutic targets.

**Statement of significance:** This study leverages advanced biomaterials and biofabrication techniques to engineer physiologically relevant, patient-specific, and sex-matched models of pulmonary fibrosis, addressing the critical need for pre-clinical therapeutic drug screening platforms. These human 3D lung models successfully replicated key features of fibrotic lung tissue. Tuning microenvironmental stiffness of 3D PEG-NB hydrogels to match fibrotic lung values and exposing human iATII cells and fibroblasts to pro-inflammatory biochemical cues recreated hallmark characteristics of *in vivo* fibrosis pathogenesis, including epithelial differentiation and loss, as well as fibroblast activation. The utility of these models was further validated by demonstrating responsiveness to Nintedanib, a clinically available treatment for IPF. These findings highlight the transformative potential of well-defined biomaterial-based 3D models for elucidating complex disease mechanisms and accelerating therapeutic drug discovery for chronic pulmonary diseases like idiopathic pulmonary fibrosis.

**Graphical abstract:** 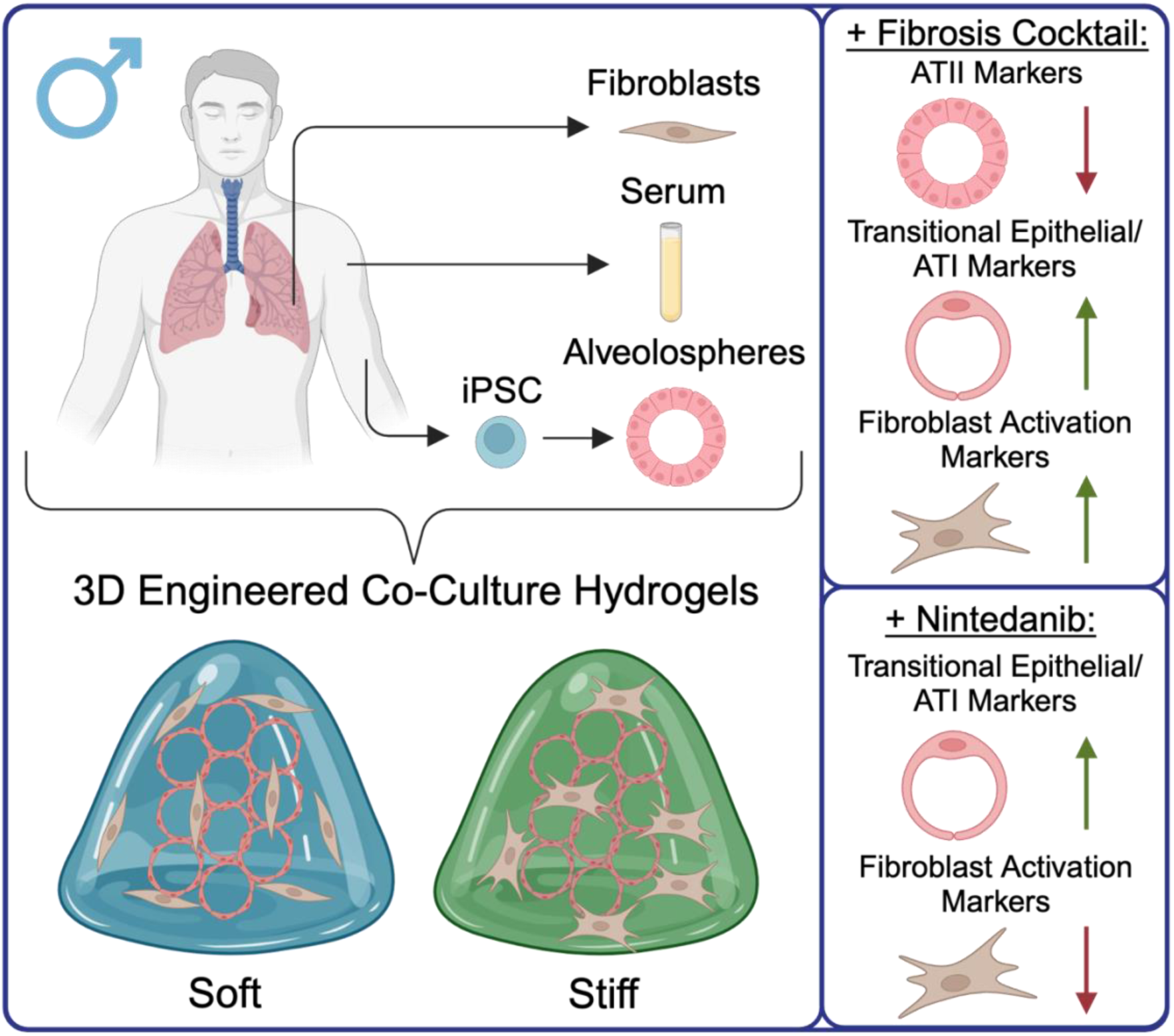

## Introduction

Idiopathic pulmonary fibrosis (IPF) is an incurable respiratory disease that results in lung tissue scarring and progressive respiratory failure [1]. Hallmarks of the disease include epithelial cell dysfunction, fibroblast activation, excessive extracellular matrix (ECM) deposition, and thus disrupted gas exchange [2–5]. The ECM provides structural support to the lungs and undergoes continuous remodeling, creating a dynamic milieu rich in biophysical and biochemical cues. Comprised of over 150 different types of proteins, enzymes, growth factors, and proteoglycans [5], the ECM plays a crucial role in lung homeostasis and disease pathogenesis. In healthy lung tissue, stiffness typically ranges between 1-5 kPa, whereas fibrotic lung tissue stiffness often exceeds 10 kPa [6, 7]. Strong evidence indicates that cell-matrix interactions are key drivers of fibrosis progression, yet the mechanisms underlying these responses are not fully elucidated [5–11]. Due to the complexity of the lung extracellular microenvironment, fully replicating *in vivo* conditions remains a significant challenge. Consequently, many researchers use reductionist models to investigate specific cellular and molecular interactions within a more controlled setting [7, 10, 12–14]. While IPF remains idiopathic, there is growing evidence that genetic and environmental risk factors [15, 16], including older age [17, 18], history of smoking or exposure to airborne hazards [19, 20], and male sex [17, 21], predispose individuals to the disease. Therefore, a need remains to engineer dynamic 3D distal lung models that support the growth of alveolar epithelial and fibroblast cells together to directly investigate the interactions between these two cell types and the surrounding microenvironment.

The alveolar region of the lungs, the primary site for gas exchange, is particularly vulnerable to damage in chronic respiratory diseases. Both lung epithelial cells and fibroblasts play an important part in IPF pathophysiology. Specifically, alveolar epithelial type II (ATII) cells, a subpopulation of alveolar epithelial cells known to produce surfactant protein C (SFTPC), function as progenitor cells within the distal lung by proliferating, differentiating, and replacing lost alveolar type I epithelial (ATI) cells [22–24], which are specialized for gas exchange and pivotal to functional epithelial repair in lung tissue. Repeated alveolar injury triggers aberrant wound-healing responses in both epithelial progenitor cells and surrounding fibroblasts. In IPF, increased impairment of epithelial regeneration results in accumulation of cells stuck in the transition from ATII to ATI characterized by markers of cell-cycle arrest, downregulation of ATII markers, upregulation of ATI markers, and high expression of unique genes including keratins, claudin-4, stratifin, and genes in the transforming growth factor-β (TGF-β) pathway [25, 26]. The cell-cycle arrest of cells in transition from ATII to ATI may result in secretion of chemokines and cytokines such as TGF-β that activate nearby fibroblasts and recruit profibrotic macrophages [25–28]. Persistence and accumulation of transitional alveolar epithelial cells have been strongly linked to disease initiation and progression, highlighting dysregulated epithelial repair as a critical area of IPF research [26, 29–31]

Primary human cells are widely used in lung models to better replicate human-specific cellular and molecular processes. However, primary ATII cells rapidly differentiate to ATI cells within days, which leads to heterogenous cell populations after approximately one week [32, 33]. To overcome this limitation and reduce confounding cellular variables, researchers have increasingly used induced pluripotent stem cell (iPSC)-derived ATII (iATII) cells that can retain a progenitor cell phenotype for months in culture [34, 35]. Traditional *in vitro* models in pulmonary regenerative medicine often only consider one cell type. Many models rely on culturing cells on substrates with supraphysiological stiffnesses that do not match lung tissue (e.g., tissue culture plastic), or neglect to investigate the three-dimensional (3D) interaction between cells and the microenvironment. Extensive evidence demonstrates that 3D culture systems more accurately mimic *in vivo* conditions, preserving cellular physiology and molecular characteristics while enhancing translational relevance [32, 34–37]. When experiments do maintain a 3D microenvironment, most protocols for iPSC differentiation and organoid culture rely almost exclusively on animal-derived materials, such as Matrigel. These materials do not provide control over mechanical properties [6, 38], geometric cues [10], or biochemical composition [39] – factors that all profoundly impact stem cell fate *in vivo* [40, 41].

Poly(ethylene glycol) norbornene (PEG-NB) hydrogels provide a versatile platform for culturing cells within well-defined 3D biomaterials, offering precise control over mechanical properties and biochemical cues in the extracellular microenvironment [8, 42]. While engineered hydrogel biomaterials are widely used across many fields, these cell culture platforms are currently underutilized in pulmonary research [8, 43]. Only a few studies have used PEG-based hydrogels to model the lung microenvironment [10, 44, 45]. To facilitate dynamic remodeling in 3D, peptide sequences degradable by cell-secreted matrix metalloproteinase (MMP) enzymes are commonly incorporated into these hydrogels [46–48]. Combining PEG-NB hydrogels with patient-derived cells improves physiological relevance of *in vitro* lung models, enables investigation of cell-cell and cell-matrix interactions driving fibrosis initiation and progression, and may accelerate the identification and validation of therapeutic targets. Drug development for IPF is particularly challenging as approximately 90% of preclinical candidates fail to demonstrate clinical efficacy in human trials [49]. Given the high cost, time-intensive nature, and stringent regulatory requirements of new drug approvals, there remains a critical need for improved *in vitro* models that expedite drug discovery.

Here we present an engineered 3D lung model that mimics important aspects of distal lung tissue. iATII spheroids were magnetically aggregated together and embedded within PEG-NB hydrogels containing pulmonary fibroblasts, replicating the acinar structure and cellular spatial arrangement found within the alveoli [44, 50]. This co-culture platform provided an environment that facilitated epithelial-fibroblast interactions [10]. Soft (elastic modulus (E) = 5.06 ± 0.33 kPa) and stiff (E = 18.90 ± 3.19 kPa) hydrogel microenvironments supported cell viability (>75%) while effectively recapitulating the mechanical properties of healthy and fibrotic lung tissue, respectively. Beyond mechanical stiffness, pro-inflammatory biochemical cues, previously described as a fibrosis cocktail, were supplemented into the culture medium to induce epithelial injury and subsequent fibroblast activation [13, 51, 52]. Gene expression analyses revealed that epithelial and fibroblast responses within stiff hydrogels exposed to the fibrosis cocktail closely matched trends measured in pulmonary fibrosis patient tissues. To further validate the model as a viable platform for pre-clinical therapeutic drug screening, Nintedanib, a Food and Drug Administration (FDA)-approved anti-fibrotic drug was tested. This treatment downregulated multiple fibroblast markers and upregulated transitional and ATI markers, indicating a possible recovery in epithelial repair and a decrease in fibrotic phenotypes. These findings underscore the potential of this human co-culture model for studying cell-cell and cell-matrix interactions, as well as its utility in drug screening applications.

## Materials and methods

### 2.1 PEG-NB synthesis

As previously published, terminal residue conjugation of an eight-arm, 10 kg/mol PEG-hydroxyl macromer (PEG-OH) produced norbornene functionalized end groups [10, 53]. In brief, PEG-OH (5 g, JenKem Technology) was lyophilized (∼ 0.1 mBar, ∼ - 80°C) and subsequently dissolved in ∼35 mL anhydrous dichloromethane (DCM; Sigma-Aldrich, cat. #270997-1L) under moisture-free conditions in a flame-dried Schlenk flask. 4-Dimethylaminopyridine (DMAP; 0.24 g, .002 mol, Acros Organics, cat. #148270050) was added to the flask and pyridine (1.61 mL, 0.02 mol, Sigma Aldrich 494410) was injected dropwise into the reaction mixture. Separately in a second flame-dried Schlenk flask, N,N’-Dicyclohexylcarbodiimide (DCC; 4.13 g, 0.02 mol, Fisher Scientific, cat. #AC113901000) was dissolved in anhydrous DCM, again under moisture-free conditions. To this flask, norbornene-2-carboxylic acid (4.9 mL, 0.04 mol, Acros Organics, cat. #453300250) was added in a dropwise manner. After 30 minutes of stirring at room temperature, the reaction mixture was filtered through Celite 545 (EMD Millipore, cat. #CX0574-1). Then, the filtrate was added to the first flask and left to react for 48 h (while protected from light). A series of wash steps using 5% sodium bicarbonate, saturated brine (∼40 grams of sodium chloride in 100 mL deionized water), and deionized water removed undesired byproducts. Each time the reacted polymer was mixed and left to separate out into two phases for approximately 5 minutes using a separatory funnel. Anhydrous magnesium sulfate (Fisher Scientific, cat. #M65-500) was added to the organic elute to remove excess water and then filtered out using filter paper (Cytiva, cat. #1002-150). The organic product was precipitated with cold diethyl ether (Fisher Scientific, cat. #E1384) and then concentrated with a rotary evaporator.

Following a 4°C overnight incubation, the diethyl ether was removed using vacuum filtration. The precipitate was vacuum dried at room temperature in a desiccator overnight, again protected from light. Dialysis with the precipitate occurred at room temperature over 72 h, where the 3.5 L of deionized water was changed four times daily. After dialysis, the product was collected and lyophilized (∼ 0.1 mBar, ∼ -80°C) to obtain a solid white powder.

Nuclear magnetic resonance (NMR) spectroscopy confirmed the end-group functionalization and purity of the PEG-NB. A Bruker DPX-400 FT NMR spectrometer was used to collect the ^1^H NMR spectrum of the product using 184 scans and a 2.5 s relaxation time. Only synthesis products above 90% functionalization were used throughout these experiments (Supplemental Fig. 1), and chemical shifts for protons (^1^H) were recorded relative to deuterochloroform as parts per million (ppm).

### 2.2 iATII cell culture and magnetic labeling

iATIIs, generously provided by the Kotton Laboratory (Center for Regenerative Medicine, Boston University) and commercially known as BU3 NGST cells (RRID: CVCL_WN82), containing thyroid transcription factor <u>N</u>KX2 homeobox 1 <u>G</u>reen fluorescent protein (NKX2-1^GFP^) and <u>S</u>urfactant protein C td<u>T</u>omato (SFTPC^tdTomato^) reporters, were maintained in 40 μL of 8 mg/mL growth factor reduced Matrigel (Corning, cat. #356231) and CK + DCI medium as previously described [34, 35]. During routine passaging, 0.05% Trypsin-EDTA (∼15 min, Gibco, cat. #25-300-062) was used to dissociate iATIIs back into a single cell state [34, 35]. Nanoshuttle (1μL per every 10,000 cells, Greiner Bio-One, cat. #657846) was then added to a proportion of the iATIIs so that the cells could be magnetically aggregated a few days later. This was done by pipetting the cells and Nanoshuttle up and down gently until visibly homogenous (1-2x) and then centrifuging at 300 x g for 5 minutes at 4°C. This process was repeated an additional two times before the iATII pellet was resuspended in 40 μL of 8 mg/mL Matrigel and standard passaging protocols were resumed [34, 35]. Nanoshuttle is a nanoparticle assembly that consists of gold, iron oxide, and poly-L-lysine. This mixture enables the Nanoshuttle beads to attach to the cell membranes electrostatically. iATIIs with Nanoshuttle were left to grow into small alveolospheres for 4-5 days prior to use in experiments.

### 2.3 Fibroblast cell culture

Frozen vials of patient-specific human lung fibroblasts (Table 1) were thawed and expanded at 37°C and 5% CO_2_ in T75 flasks containing growth medium (Dulbecco’s Modified Eagle Medium (DMEM), 10% v/v charcoal-stripped fetal bovine serum (CS-FBS, Table 1), and 1% v/v penicillin/streptomycin). All fibroblasts used in experiments were seeded between passages two and seven.

**Table 1.**
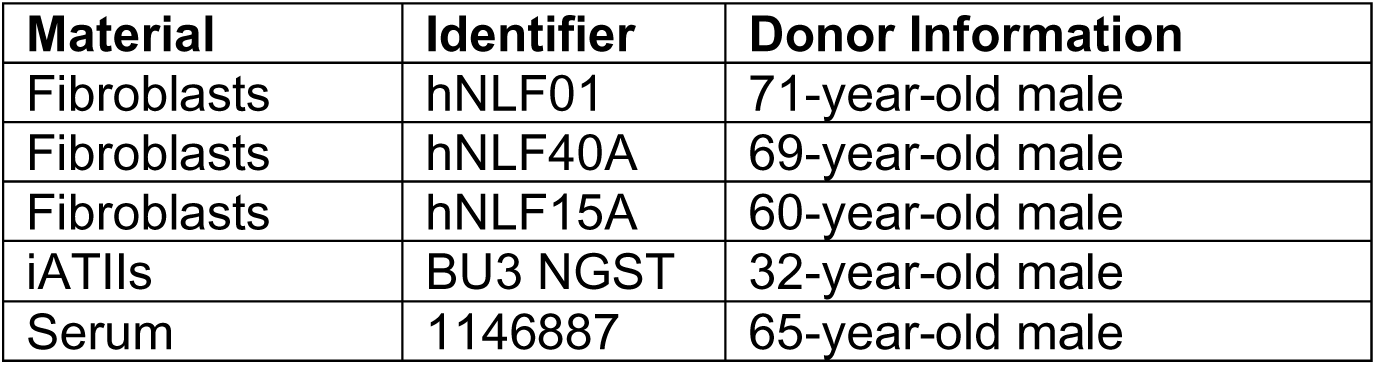
Human cell and serum information.

### 2.4 Preparation of the embedding hydrogel

The initial PEG-NB weight percent determined whether the embedding hydrogel corresponded to a soft (5.25 wt%) or stiff (7.75 wt%) formulation, with a 0.7 ratio of thiols to norbornenes. A matrix metalloproteinase-9 (MMP9)-degradable peptide (Ac-GCRD-VPLSLYSG-DRCG-NH2, GL Biochem) was used as a crosslinker and both fibronectin (CGRGDS, 2 mM, GL Biochem) and laminin (CGYIGSR, 2 mM, GL Biochem) mimetic peptides, as well as 2 mg/ml Laminin/Entactin (Corning, cat. #354259) were incorporated into the formulation to enhance cell adhesion. Lithium phenyl-2,4,6-trimethylbenzoylphosphinate (LAP, 1.1 mM) acted as the photoinitiator. Cell culture medium (CK + DCI) was used to reconstitute the PEG-NB, MMP9- degradable peptide crosslinker, CGRGDS, and CGYIGSR. The individual volumes of each component were mixed, and the overall pH of the final hydrogel precursor solutions (soft: 5.25 wt% PEG-NB, 13.29 mM MMP9, 2 mM CGRGDS, 2 mM CGYIGSR, 2 mg/mL laminin/entactin, and 1.1 mM LAP; stiff: 7.75 wt% PEG-NB, 19.62 mM MMP9, 2 mM CGRGDS, 2 mM CGYIGSR, 2 mg/mL laminin/entactin, and 1.1 mM LAP) were adjusted to pH∼7-8 as needed.

### 2.5 Rheological characterization of acellular hydrogels

Hydrogels for rheological evaluation were prepared by pipetting 40 μL of final hydrogel precursor solution between two glass slides covered in parafilm and separated by a 1 mm gasket. After ultraviolet (UV) light exposure at 365 nm with 10 mW cm^−2^ intensity (Omnicure, Lumen Dynamics) for 5 min, these samples polymerized. Hydrogels were then swollen in phosphate buffered saline (PBS) overnight prior to characterization. The elastic modulus of the hydrogels (e.g., stiffness) was measured with an 8-mm parallel plate geometry on a Discovery HR2 rheometer (TA Instruments) as previously described [14, 53]. In brief, a hydrogel was placed onto the Peltier plate (37℃) and the geometry was lowered until it was in contact with the hydrogel surface and an axial force of 0.03 N was registered. The storage modulus (G’) plateau was determined by measuring the storage modulus at different 5% increments of compression until a maximum was reached [10, 53]. The storage modulus plateau for the soft hydrogels occurred at 25% compression, and 30% compression for the stiff hydrogels. The hydrogel samples then underwent a frequency oscillation with logarithmic sweep of frequencies (1-100 rad s^-1^) and 1% strain. From here, the elastic modulus (E) was calculated under the assumption that the hydrogels were incompressible and exhibited bulk-elastic characteristics with a Poisson ratio of 0.5 [7, 54–56].

### 2.6 Formation of 3D acinar structures

Both enzymatic (2 mg/mL dispase; Thermo Fisher Scientific, cat. #17105-041) and manual pipetting were used to free the iATII alveolospheres from Matrigel constructs over 30 minutes. The alveolospheres were then rinsed in DMEM and pelleted, using centrifugation at 300 x g for 5 minutes and 4°C. This washing step was completed a total of 3x to ensure the enzyme was completely removed. Following the last spin, the alveolospheres were resuspended in CK + DCI medium (Supplemental Table S1; [34, 35]) supplemented with 10 μm Y-27632 (Tocris, cat. #1254), counted, and then transferred into 24-well cell-repellent plates (Greiner Bio-One, cat. #662970) at a concentration of 400 spheres and 250 μL of CK + DCI + 10 μm Y-27632 medium per well. Magnetic levitation drives (Bio-Assembler, Greiner Bio-One, cat. # 662840) were added to the plates and then transferred onto an orbital shaker (∼60 rpm) within a cell culture incubator (37°C, 5% CO_2_) to facilitate aggregation of alveolospheres into 3D acinar structures. After 3 hours, the magnetic levitation drives were removed and replaced with magnetic concentrating drives (Greiner Bio-One, cat. # 662840) and the epithelial cells were allowed to settle over approximately 5 minutes. At this point, medium was removed manually from each well and a hydrophobic pen (Vector Laboratories, cat. #H-4000) was used to draw a circle around the cells. This barrier ensured that the alveolospheres were completely encapsulated in the hydrogel precursor solution prepared in the subsequent steps. It was important to minimize the amount of time the iATIIs were left without medium to ensure high viability.

### 2.7 Hydrogel embedding of 3D acinar structures

Just prior to embedding, fibroblasts were dissociated into single cell with trypsin, assessed for viability with Trypan Blue, and counted on a hemocytometer. Fibroblasts were pelleted and then resuspended in the precursor hydrogel solution, so the final concentration was 1,000 fibroblasts/μL. A total of 40 μL of embedding hydrogel precursor solution containing fibroblasts (40,000 fibroblasts total per sample) was added directly on top of the exposed alveolosphere aggregate. Five minutes of UV light exposure (365 nm, 10 mW cm^−2^, Omnicure, Lumen Dynamics) polymerized the hydrogels. Prior work has established that there are no significant differences in cell viability and transcriptome when cells are exposed to 365 nm light for this length of time [57]. Afterwards, these samples were carefully transferred into new 24 well plate wells that contained CK + DCI medium [34, 35] that was supplemented with 10 μm Y-27632 and 1% serum from a 65-year-old male patient (Table 1) for 48 h (37°C and 5% CO_2_).

### 2.8 Fibrosis cocktail exposure

Samples were maintained at 37°C with 5% CO_2_ in CK + DCI medium [34, 35] supplemented with 1% serum from a 65-year-old male patient (Table 1). For fibrotic activation experiments, samples were either exposed to a fibrosis cocktail (FC) or vehicle control (VC). The fibrosis cocktail contained 5 ng/ml recombinant transforming growth factor beta (TGF-𝛽; PeproTech, cat. #100-21), 10 ng/ml platelet-derived growth factor AB (PDGF-AB; Thermo Scientific, cat. #PHG0134), and 5 μM 1-Oleoyl Lysophosphatidic Acid (LPA; Cayman Chemical Company, cat. #62215) [13, 51, 58, 59]. Dosing began on day 2 and continued until day 8, where each well was replenished with medium (1 mL/well) containing the FC or VC (PBS supplemented with 0.1% bovine serum albumin (BSA)) every 48 h.

### 2.9 Live-dead and immunofluorescence staining

Commercially available human pulmonary fibroblasts (HPFs) were used for all viability studies (passage 2-7). A ReadyProbes Cell Viability Imaging Kit (Thermo Fischer Scientific, R37609) quantified the number of live cells in each construct.

Samples treated with FC or VC were collected for imaging after 2, 4, 6, or 8 days. Prior to imaging, 1 drop of NucBlue (nuclei) and 1 drop of NucGreen (dead) was added to each 1 mL of cell culture medium to make a staining medium. Samples were transferred into 24-well plate wells containing 300 μL of staining medium and on an orbital shaker for 1 h (37°C, 5% CO_2_). Afterwards, samples were transferred onto a glass slide and covered in PBS to maintain hydration during imaging. A hydrophobic pen was used to confine the PBS to the sample area.

The fluorescently stained samples were imaged on an Olympus CKX53 upright microscope adapted for fluorescent capabilities with DAPI and FITC filters. Six random points in the construct were imaged at 4x using 100 ms exposures for the DAPI (nuclei) channel,10-20 ms for the FITC (dead) channel, and 100 ms for the TRITC (SFTPC) channel. Exposures were kept constant each day of imaging and between samples that were directly compared. Images were post-processed and analyzed with ImageJ software (NIH). Total cell viability was quantified using Equation 1.

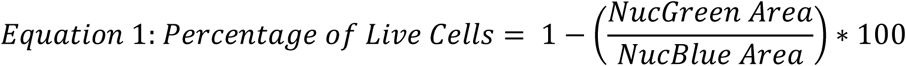

Also, to visually assess where fibroblasts were within the hydrogel relative to the iATIIs, CellTracker Green CMFDA (10 μM, Thermo Fisher, cat. #C7025) was used for non-viability related samples. This fluorescently tagged the fibroblasts green after incubating the CellTracker Green CMFDA on the cells for 45 min in serum free medium. Following, the fibroblasts were dissociated with trypsin, embedded, and imaged while in culture.

### 2.10 Magnetic-activated cell sorting (MACS)

Epithelial cells and fibroblasts were purified out of co-culture using magnetic column isolation based on the expression of EpCAM (CD326), which is a cell surface marker that is found on ATII, transitional epithelial, and ATI cells [10, 60, 61]. On days 6 and 8, an enzymatic digestion solution containing 5 mg/mL dispase and 1 mg/mL elastase (Worthington Biochemical, cat. #LS002292) was prepared fresh in DMEM. Old medium was removed from each well and replaced with 700 μL of the enzymatic digestion solution. Both enzymatic and manual pipetting were used to free the cells from hydrogel constructs, which took up to 1 h. The dispase and elastase enzymes targeted the MMP9-degradable crosslinker sequence and helped facilitate degradation. Four to six samples of the same experimental group were pooled together to form each technical replicate. Once hydrogel degradation occurred, the solution of iATIIs and fibroblasts was transferred into 15 mL test tubes, diluted 1:1 with DMEM, and then centrifuged at 300 x g for 5 minutes and 4°C to create a cell pellet. Trypsin-EDTA (0.05%; Gibco, cat. #25-300-062) was added to the cell pellets and then transferred into 6-well plates to allow for the alveolospheres to dissociate into single cells for approximately 16 min, with manual pipetting (∼2x) at the halfway timepoint. To deactivate the trypsin, a 10% v/v CS-FBS in DMEM medium was then added to the samples and then centrifuged for 5 mins at 300 x g for 5 minutes and 4°C. Next, the supernatant was manually discarded, and the cell pellets were resuspended in 10 μL of anti-CD326 (epithelial cell adhesion molecule (EpCAM)) Microbeads (Miltenyi Biotec, cat. #130-061-101) and 70 μL of buffer that consisted of 2 mM ethylenediaminetetraacetic acid (EDTA; ThermoFisher, cat. #AM9260G) and 0.5% bovine serum albumin (BSA; Sigma-Aldrich) in PBS (PEB buffer). These samples were incubated at 4°C for 15 min to allow bead binding to cells before an additional 1 mL of PEB buffer was added to each and the test tubes were centrifuged at 300 x g for 5 minutes and 4°C. From each cell pellet, the supernatant was removed, and the cells were resuspended in 500 μL of PEB buffer, ensuring there were no cell clumps.

iATIIs were positively selected using a MiniMACS Separator and Starting Kit (Miltenyi Biotec, cat. #130-090-312) set up and MS columns (Miltenyi Biotec, cat. #130- 042-201) according to the manufacturer’s protocol. Briefly, the MS columns were primed by passing 500 μL of PEB buffer through the columns. Then, a test tube was placed beneath the column and the cells in 500 μL of PEB buffer was passed through to allow the EpCAM^+^ cells bound to beads to be trapped in the magnetic field. An additional two rinse steps of 500 μL of PEB were done to ensure all EpCAM^−^ fibroblasts were collected in the test tube below. Lastly, a new test tube was placed beneath the column at this point, and 1 mL of PEB buffer was added to the column. The column was then removed from the magnet holder, and the plunger was compressed to elute the positively selected EpCAM^+^ cells into the second test tube. Each respective test tube containing cells was centrifuged one final time at 300 x g for 5 minutes and 4°C.

### 2.11 RNA isolation and cDNA synthesis

Immediately following MACS isolations, the supernatant was removed from the iATII and fibroblast cell pellets, and 300 μL of TRIzol Reagent (Fisher Scientific, cat. #15-596-026) was added to each test tube. The samples were pipetted and briefly vortexed before being stored at -20°C for up to one month. After samples were thawed, the sample volume was transferred into a 1.5 mL Eppendorf tube. 100 μL of 1-bromo-3- chloropropane (BCP; Fisher Scientific, cat. #NC9551474) was added to each Eppendorf tube. Next, each Eppendorf tube was briefly vortexed, incubated at room temperature for 5 min, and incubated on ice for an additional 5 min. At this time, the Eppendorf tubes were centrifuged at 12,000 x g for 15 min, so the clear aqueous layer of the sample could be transferred into a separate RNAse-free 1.5 mL microcentrifuge tube. An equal amount of 100% ethanol (EtOH) was added to the clear RNA layer volume and the two were briefly vortexed. Up to 700 μL of total volume was transferred into a RNeasy Plus

Micro Kit column (Qiagen, cat. #74034) and purified according to the manufacturer’s instructions. RNA quantity and purity, as assessed by the ratio of 260 nm and 280 nm (A_260_/A_280_) absorbance readings, were measured using a BioTek plate reader and a Take3 Micro-Volume Plate. The isolated RNA was then converted into cDNA using a high-capacity cDNA Reverse Transcription Kit (Applied Biosystems, cat. #4368814) according to the manufacturer’s protocol.

### 2.12 Assessment of cell specific gene expression

Reverse transcription-quantitative polymerase chain reaction (RT-qPCR) assessed gene expression for a variety of different ATII (*SFTPC, LAMP3*), ATII-ATI (*KRT17, CLDN4*), ATI (*PDPN, AQP5*), aberrant basaloid cell (*FN1*), and fibroblast activation (*COL1A1, FN1, CTGF, CTHRC1, LTBP2*) markers. iTaq Universal SYBR Green Supermix (Bio-Rad, cat. #1725121) and a CFX Opus 96 (Bio-Rad) were used for all experiments. Fibroblast gene expression was normalized to ribosomal protein L30 (*RPL30*), which served as the housekeeping gene, whereas epithelial gene expression was normalized to ribosomal protein S18 (*RPS18*). All human primers were acquired from Integrated DNA Technologies (Supplemental Table S2). All Ct values were natural log transformed to normalize the data for statistical analyses [62], and then relative gene expression was calculated using a 2^−Δ𝐶𝑡^ approach. After statistical analyses were computed, values were untransformed (by taking the exponent of the natural logged value) and presented in Fig.s.

### 2.13 Validating response to anti-fibrotic drug treatment

To narrow down conditions for therapeutic drug testing, the influence of the age of patient fibroblasts, substrate elastic modulus, exposure, and time on either epithelial gene expression or fibroblast activation gene expression were investigated with a design of experiment (DOE) approach using JMP software (Pro 18 Version, SAS). The resulting least-squares regression model identified the stiff hydrogel formulation as the most fibrotic microenvironment. Therefore, all subsequent drug studies were only conducted with patient specific fibroblast cells in stiff hydrogels and gene expression was assessed at the day 8 timepoint. For drug treatment, 10 μM of Nintedanib (Tocris, cat. #7049) in dimethyl sulfoxide (DMSO) was supplemented into the cell culture medium (CK + DCI + 1% male serum) simultaneously with the FC components starting on day 4 and replenished every 48 h at the subsequent medium changes. Meanwhile, samples that were kept in cell culture medium (CK + DCI + 1% male serum) with only the FC components served as controls.

### 2.14 Statistical methods

For each viability timepoint, images were acquired from six individual hydrogels (n=6 technical replicates per experimental condition). Unpaired t-tests (GraphPad Prism) were used to calculate statistical significance for day 2 viability results, whereas two-way ANOVAs followed by Tukey’s honest statistical difference (HSD) tests (GraphPad Prism) were applied to compare VC and FC results on days 4, 6, and 8.

Similarly, for rheological measurements, six separate hydrogels (n=6 technical replicates per stiffness) were measured. An unpaired t-test was used to calculate statistical significance for these data sets (GraphPad Prism). For all MACS isolations, 4-6 samples of the same experimental group were pooled together to form each biological replicate. In total, the RT-qPCR results presented in Fig.s 4 and 5 considered three biological replicates (3 separate patient specific fibroblast lines, N=3). Each relative gene expression value was then entered back into the DOE and a standard least squares model was applied to fit the model and identify best fit lines. The JMP software identified a multi-factorial design and approached the statistical analysis similarly to a three-way ANOVA. For the drug testing, 6 hydrogels were pooled together, and RT-qPCR results were collected using the 3 separate patient specific fibroblast lines (N=3). Statistical significance was determined with paired t-tests for these results.

## Results

### 3.1 PEG-NB hydrogels recapitulated key aspects of fibrotic tissue

To engineer a cellular microenvironment that recreated fibrosis progression *in vitro*, iATIIs and fibroblasts were embedded in well-defined, tunable stiffness 3D PEG-NB hydrogels (Fig. 1A). These hydrogel formulations were comprised of an eight-arm, 10 kg/mol PEG macromer that was 93% functionalized with norbornene end groups (Supplemental Fig. S1), an MMP9-degradable peptide crosslinker, a fibronectin mimetic peptide (CGRGDS), a laminin mimetic peptide (CGYIGSR), and entrapped laminin/entactin protein complex (Fig. 1A). Both the pendant peptides and the laminin/entactin protein complex facilitated increased cellular adhesion to hydrogels.

**Fig. 1.**
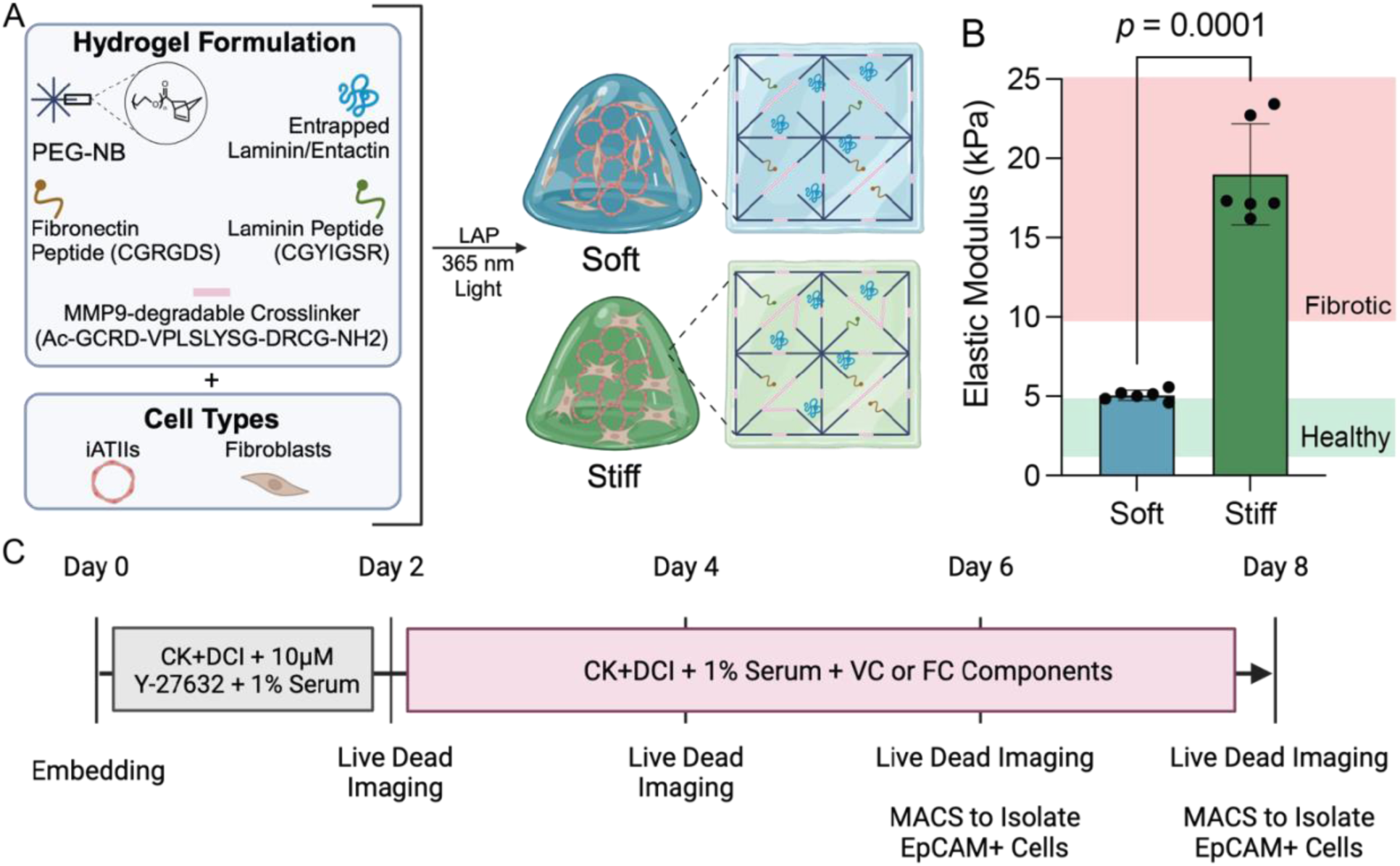
Engineered hydrogels supported iATII-fibroblast co-cultures with stiffnesses tuned to match healthy and fibrotic lung tissue. (A) Cells were embedded in 3D arrangements mimicking alveolar micro-architecture within soft or stiff hydrogel formulations that contained PEG-NB, an MMP9-degradable crosslinker, CGRGDS, CGYIGSR, and entrapped laminin/entactin. (B) Rheological measurements for the average elastic modulus (E) of soft (E = 5.06 ± 0.33 kPa, n=6) and stiff hydrogels (E = 18.99 ± 3.19 kPa, n=6). Soft and stiff hydrogel formulations fall within the ranges for healthy (green region) and fibrotic (red region) lung tissue stiffness. Columns represent mean ± SD. Symbols represent technical replicates. Statistical significance was determined by an unpaired t-test. (C) Schematic representation of the experimental timeline and outputs.

Modification of the PEG weight percent varied the elastic modulus (e.g., stiffness) to produce hydrogels with discrete stiffnesses. Soft hydrogels demonstrated an elastic modulus of 5.06 ± 0.33 kPa, while stiff hydrogels exhibited an elastic modulus of 18.90 ± 3.19 kPa, effectively matching the mechanical properties of healthy and fibrotic lung tissue [6, 7], respectively (Fig. 1B). iPSC-derived iATIIs and fibroblasts were arranged within these hydrogels to replicate 3D lung micro-architecture. First, Nanoshuttle, magnetic nanoparticles attached to poly-l-lysine, was attached to iATII cell membranes through electrostatic attraction. The magnetized iATII alveolospheres were aggregated together within a magnetic field using bioassembler magnetic drives (Greiner) to form acinar like structures. Then, fibroblasts were distributed throughout the embedding hydrogel precursor solution and added on top of the iATII aggregates. As a result, the samples consisted of an epithelial core surrounded by fibroblasts, which enabled epithelial-fibroblast crosstalk within 3D lung models. Fig. 1C describes the different experimental outputs and mediums that were used over the course of the 8-day experimental timeline. Samples were kept in CK+DCI medium supplemented with 1% human serum and a rock inhibitor for the first 48 hours. On day 2, the medium was then switched to CK+DCI medium supplemented with 1% human serum and either the fibrosis cocktail (FC) or vehicle control (VC) components. This medium was replenished every 48 h until day 8. Cell viability was assessed on days 2, 4, 6, and 8 while magnetic column isolations to separate epithelial and fibroblast cellular subpopulations occurred on days 6 and 8 (Fig. 1C).

### 3.2 Cells maintained high viability in 3D lung models

Total cellular viability in 3D distal lung models was measured with a ReadyProbes Cell Viability Imaging Kit at days 2, 4, 6, and 8 for each of the four conditions combining two different stiffness (soft or stiff) hydrogels and two different exposures (VC or FC). Representative images showed all nuclei marked by blue fluorescence, dead cells marked by green fluorescence, and the iATIIs marked by red fluorescence from the SFTPC^tdTomato^ reporter (Fig. 2A). After the embedding process on day 2, 92.11% ± 4.71% of cells remained alive within the soft hydrogels, while 92.14% ± 14.40% of cells remained alive within the stiff hydrogels (Supplemental Fig. S2). Cell viability within the soft hydrogels measured approximately 98% at day 4, and maintained at 96% by day 8, with no differences in viability between VC and FC conditions (Fig. 2B). Stiff hydrogel cell viability was 85% at day 4 and increased slightly to 92% by day 8, indicating that even if the stiff microenvironment induced some initial cell death post-embedding, overall, these models promoted high cell viability (Fig. 2C). The results also showed that there were no statistically significant differences in cell viability between the VC and FC exposures for both soft and stiff hydrogels across all timepoints (Fig. 2C).

**Fig. 2.**
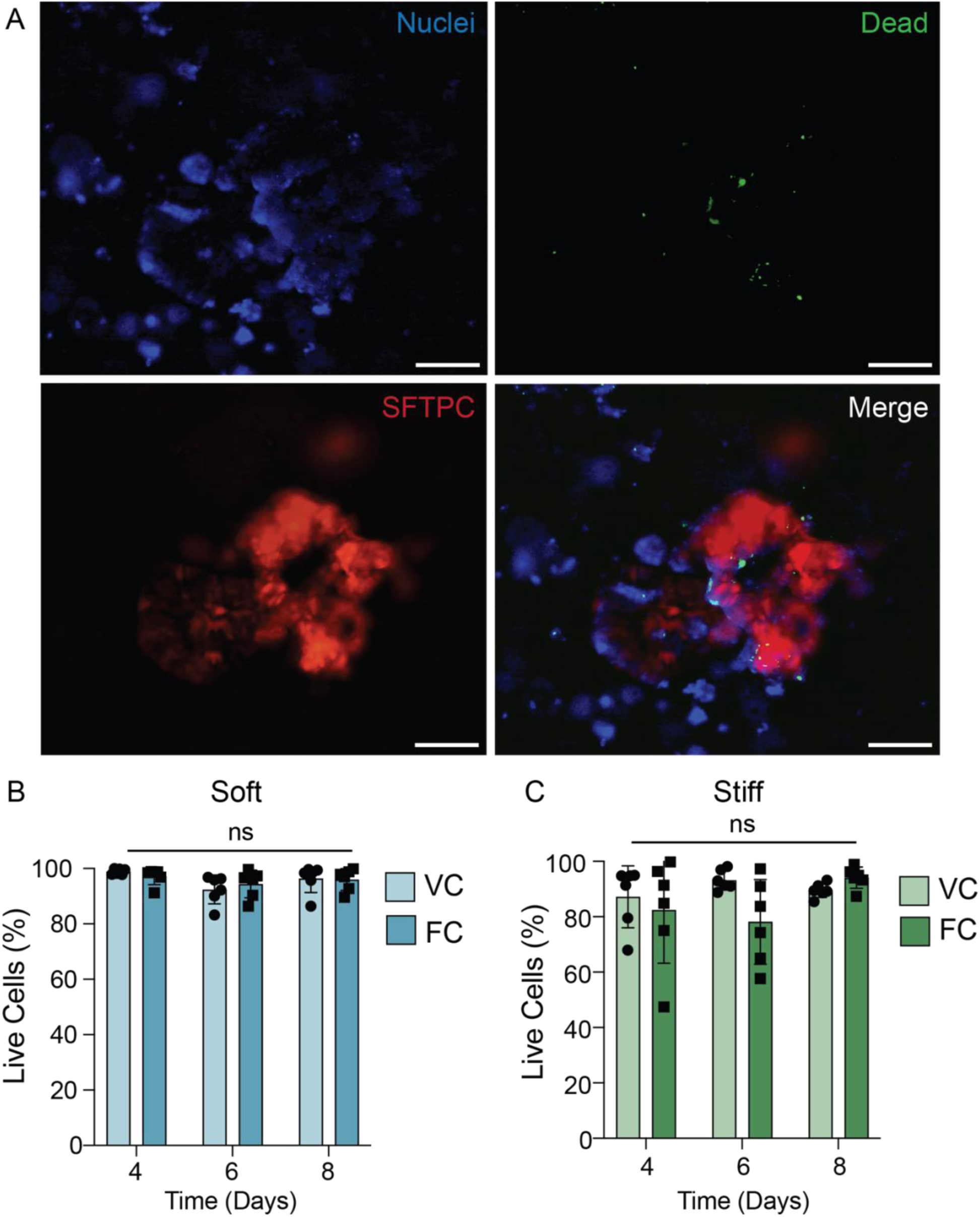
Cells maintained viability in 3D hydrogels over the 8 days in culture. (A) Representative images of soft 3D hydrogels exposed to the fibrosis cocktail with cells stained with a ReadyProbes cell viability imaging kit on day 4. Cell nuclei were stained blue, dead cells were labeled green, and SFTPC^tdTomato^ reporter expression associated with iATII cells was visualized as red fluorescence. Scale bars = 200 µm. (B) Quantification of cell viability in soft hydrogels either treated with the vehicle control (VC) or fibrosis cocktail (FC). Columns represent mean ± SD, n=6. Symbols represent technical replicates. Statistical significance was assessed by a two-way ANOVA with Tukey’s multiple comparisons test, ns = no significance. (C) Quantification of cell viability in stiff hydrogels either treated with VC or FC. Columns represent mean ± SD, n=6. Symbols represent technical replicates. Statistical significance was assessed by a two-way ANOVA with Tukey’s multiple comparisons test, ns = no significance.

### 3.3 Pro-inflammatory biochemical cues induced epithelial damage and fibroblast activation

3D lung models successfully replicated key geometric and spatial cellular characteristics of distal lung tissue. Magnetic aggregation of iATII alveolospheres formed an epithelial structure replicating alveolar architectures (Fig. 3A). Epithelial aggregates maintained robust tdTomato (red) fluorescence, and a high number of SFTPC^+^ cells at the time of embedding. To visualize the spatial arrangement of fibroblasts relative to the epithelial core, CellTracker labeled the fibroblasts green (Fig. 3B). The proximity of fibroblasts to epithelial aggregates within these 3D lung models enables cell-cell interaction as described in our previous studies [10]. After samples were in culture, FC was supplemented into the medium for half of the samples to provide pro-inflammatory cues and further induce fibrotic activation independent of microenvironmental stiffness. The FC (5 ng/ml TGF-𝛽, 10 ng/ml PDGF-AB, and 5 μM LPA) was initially added on day 2, and replenished every 48 hours until day 8. Magnetic column isolations positively selected for EpCAM^+^ cells on days 6 and 8, enabling cell- specific relative gene expression. This approach ensured ATII, ATI, and alveolar epithelial transitional cells were separated from the fibroblasts. Relative gene expression results determined by RT-qPCR were visualized within a heat map (Fig. 3C). Surfactant protein C (*SFTPC*) and lysosome-associated membrane protein 3 (*LAMP3*) served as ATII markers, keratin 17 (*KRT17*) and claudin-4 (*CLDN4*) served as ATII-ATI transitional epithelial cell markers, podoplanin (*PDPN*) and aquaporin-5 (*AQP5*) served as ATI markers. Fibroblast activation markers were collagen 1 alpha chain 1 (*COL1A1*), fibronectin (*FN1*), connective tissue growth factor (*CTGF*), collagen triple helix repeat containing 1 (*CTHRC1*), and latent TGF-β binding protein 2 (*LTBP2*). Increased microenvironmental stiffness and FC exposure were expected to downregulate ATII gene expression, and increase ATII-ATI, ATI, and fibroblast activation gene expression. Relative gene expressions followed these expected results across nearly all genes, with a notable deviation occurring with the *AQP5* gene expression (Fig. 3C). Overall, the heat map results supported the hypothesis that epithelial damage from FC exposure led to epithelial differentiation and contributed to fibroblast activation.

**Fig. 3.**
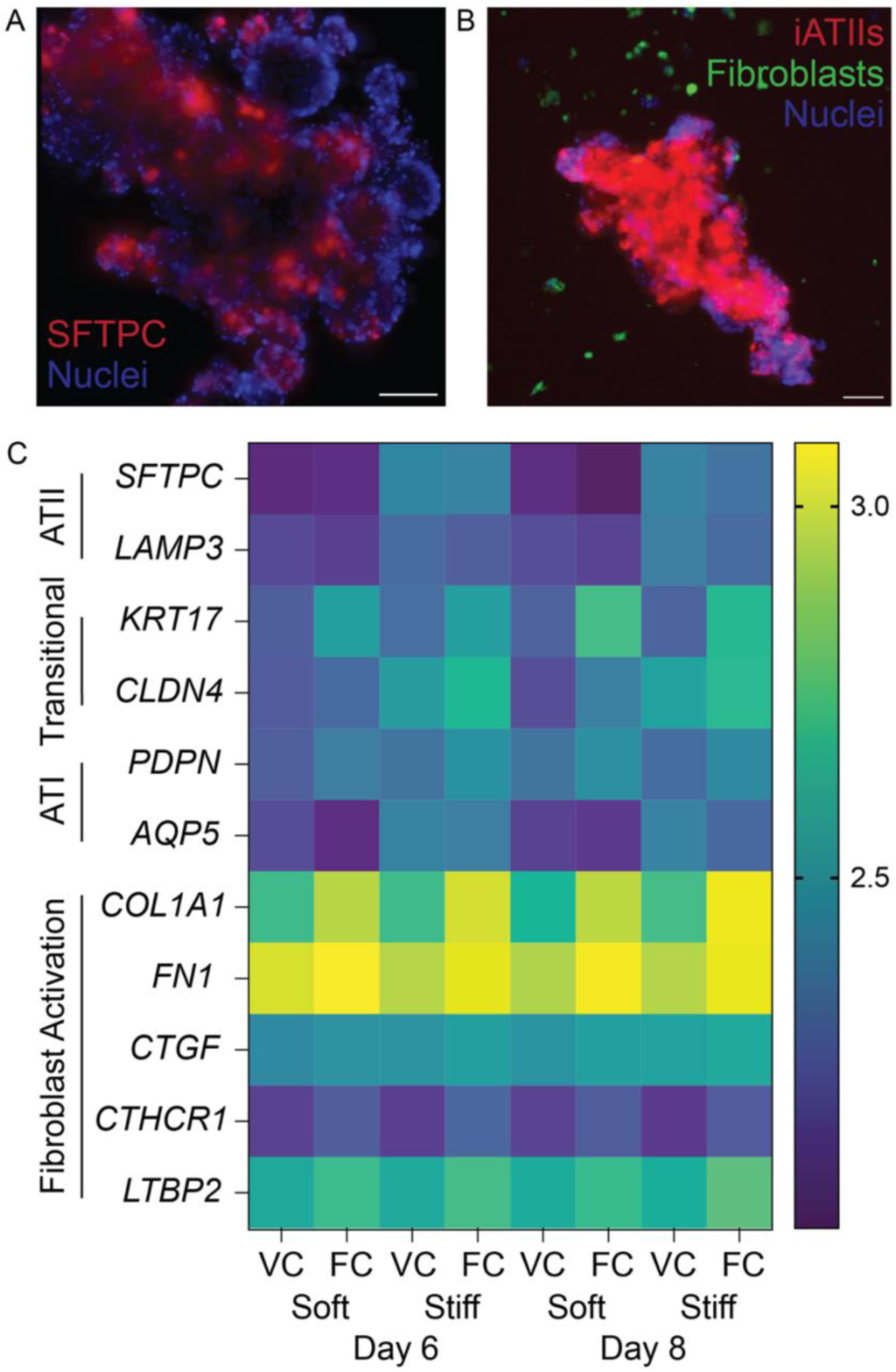
iATII-fibroblast co-culture recapitulated characteristics of fibrotic tissue. (A) Representative maximum intensity projection of a confocal image of epithelial aggregate with iATIIs expressing *SFTPC* (red), counterstained for nuclei (blue). Scale bar = 150 µm. (B) Representative maximum intensity projection of a confocal image of 3D lung models in PEG-NB hydrogel showing the spatial arrangement of iATIIs expressing *SFTPC* (red) near alveolar fibroblasts (green), counterstained for nuclei (blue). Scale bar = 100 µm. (C) Heat map showing relative gene expression of ATII, transitional epithelial, ATI, and fibroblast activation markers. Gene expression patterns in 3D lung models matched trends measured in fibrotic lung tissue (N=3 biological replicates).

### 3.4 Statistical analyses revealed the factors that created the most fibrotic conditions in 3D distal lung models

Two individual designs of experiments further statistically analyzed how the input variables of the age of patient fibroblasts, hydrogel elastic modulus, exposure (VC or FC), and day of collection for analysis influenced either epithelial or fibroblast gene expression. Primary fibroblasts were isolated from healthy male patients (Table 1) aged 60, 69, and 71 years old. The average elastic modulus measurements for the soft and stiff hydrogels were 5 kPa and 18.9 kPa. Additionally, gene expression was assessed on days 6 and 8. In response to epithelial injury, ATII cells can differentiate into ATI cells to repair damaged tissues. In pulmonary fibrosis dysregulated healing may result in the accumulation of transitional epithelial cells [25, 26]. Following *in vivo* observations, the statistical model was directed to maximize all response variables except *SFTPC* gene expression, which was minimized to replicate a fibrotic healing response. Trend lines for each output variable plotted in response to each input variable provided a visual depiction of how epithelial cells responded to each input (Fig 4A). The results revealed that individual relative gene expression varied widely based on the age of patient-derived fibroblasts, exposure, and time. However, a clearer trend emerged when assessing the input variable of elastic modulus: relative epithelial gene markers were all upregulated within the stiff hydrogels (Fig 4A). The most fibrotic condition for the epithelial cells was achieved by day 6 when the 69-year-old patient fibroblasts were embedded within stiff (18.9 kPa) hydrogels and exposed to the FC (Fig. 4A). The input variable of hydrogel elastic modulus had the greatest influence on the epithelial gene expression (p = 0.001) (Fig. 4B). The next most influential factors were the interaction between the age of patient fibroblasts and the elastic modulus (p = 0.002), exposure (p = 0.006), the age of patient fibroblasts alone (p = 0.015), the interaction between the age of patient fibroblasts with time and the elastic modulus (p = 0.019), the interaction between the time and elastic modulus (p = 0.030), time (p = 0.036), and the interaction between the age of the patient fibroblasts and exposure (p = 0.047) (Fig. 4B).

**Fig. 4.**
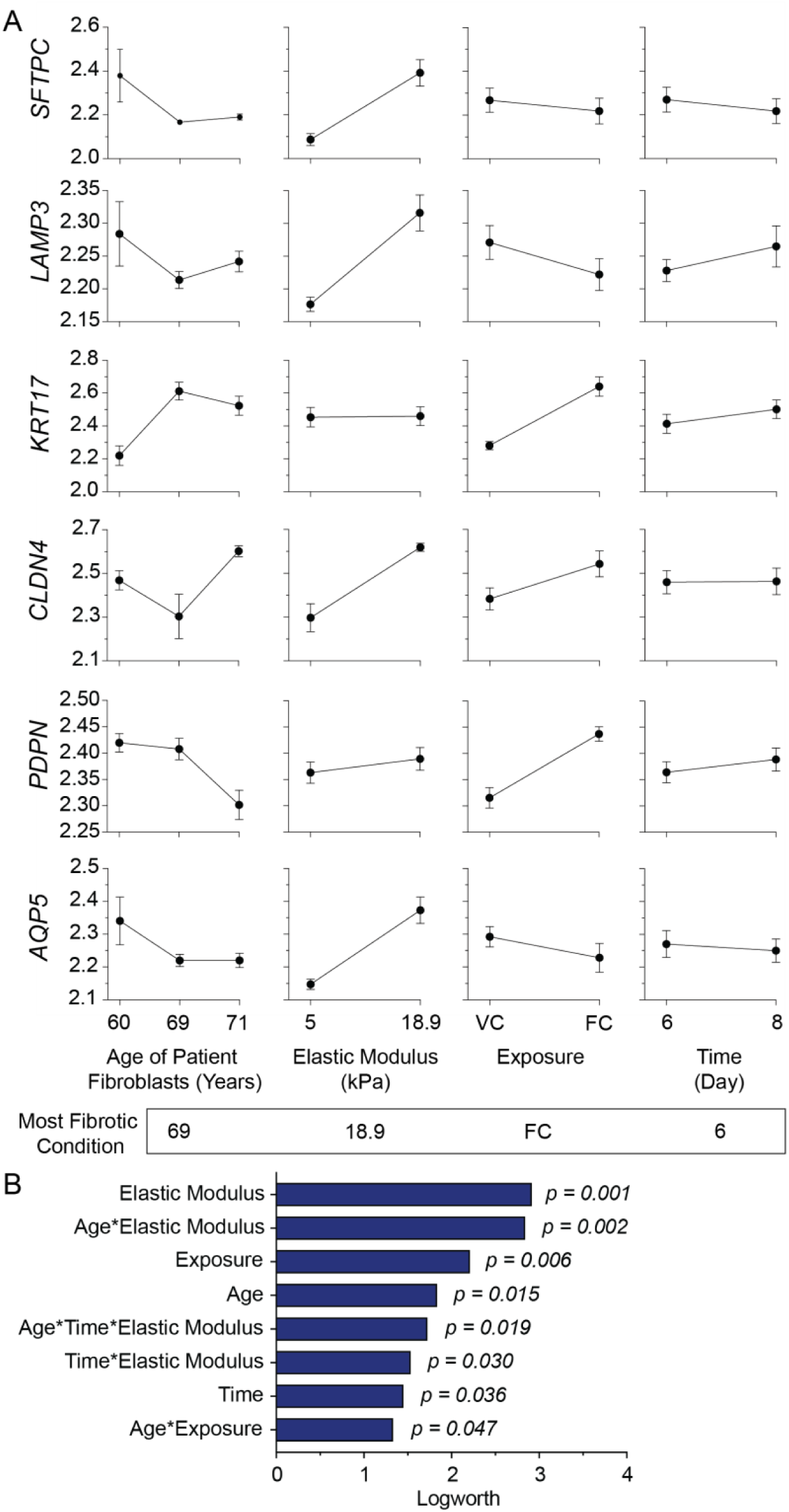
Results showed that the age of patient fibroblasts, elastic modulus, exposure, and time all significantly influenced EpCAM+ cell gene expression. (A) Trend lines from experimental data showed how *SFTPC*, *LAMP3*, *KRT17*, *CLDN4*, *PDPN*, and *AQP5* gene expression changed in relation to the age of patient fibroblasts, elastic modulus of the hydrogel, exposure, and time. Data presented as mean ± SEM. (B) The effect magnitude analysis identified all factors and interactions that were significant in influencing epithelial gene expression. These results also predicted that the most fibrotic microenvironment for EpCAM^+^ cells would be achieved by day 6 using the 69-year-old patient fibroblasts embedded in stiff (18.9 kPa) hydrogels and the fibrosis cocktail (FC).

In contrast to the epithelial results, the clearest trend for fibroblast activation emerged within the input variable of FC exposure. Relative fibroblast activation gene expression was all upregulated within the FC condition compared to VC (Fig. 5A). The fibroblast results identified that the most fibrotic condition would occur at day 8 with the 60-year-old patient fibroblasts embedded in stiff (18.9 kPa) hydrogels with FC exposure (Fig. 5A). The input variable of exposure had the greatest influence on the fibroblast gene expression (p = 0.005) (Fig. 5B). Then, the next most influential factors were the hydrogel elastic modulus (p = 0.008), time (p = 0.009), the interaction between the age of the patient fibroblasts and the elastic modulus (p = 0.010), the interaction between the age of the patient fibroblasts and exposure (p = 0.011), age (p = 0.020), and the interaction between the age of the patient fibroblasts and time (p = 0.046) (Fig. 5B).

**Fig. 5.**
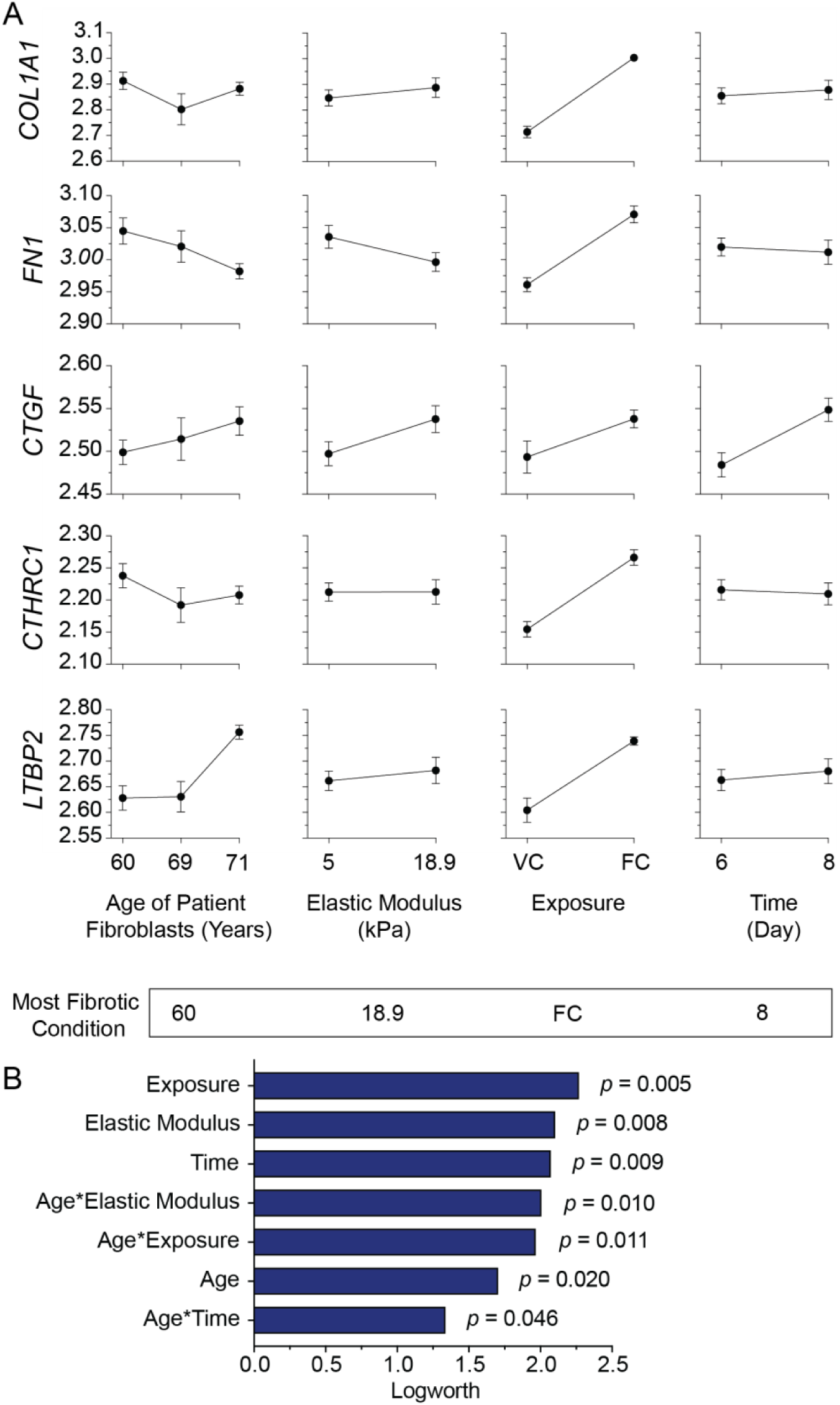
Statistical analysis results showed that the age of patient fibroblasts, elastic modulus, exposure, and time all significantly influenced fibroblast activation gene expression. (A) Trend lines from experimental data showed how *COL1A1*, *FN1*, *CTGF*, *CTHRC1*, and *LTBP2* gene expression changed in relation to the age of patient fibroblasts, elastic modulus of the hydrogel, exposure, and time. Data are presented as mean ± SEM. (B) The effect magnitude analysis identified all factors and interactions that were significant in influencing fibroblast activation gene expression. These results also predicted that the most fibrotic microenvironment for fibroblasts would be achieved by day 8 using the 60-year-old patient fibroblasts embedded in stiff (18.9 kPa) hydrogels with the FC.

### 3.5 Human 3D lung models were responsive to therapeutic drug treatment

To further evaluate 3D distal lung model responsiveness to potential drug treatments, Nintedanib was tested on co-cultured iATII cells and fibroblasts in stiff hydrogels, which represented the most fibrotic microenvironment based on the initial experiments (Fig. 4A and Fig. 5A). Cells were exposed to the FC from days 2 to 4 to induce fibrotic activation, followed by Nintedanib treatment during the remaining four days (Fig. 6A). On day 8, EpCAM^+^ cells and fibroblasts were separated using MACS, and subsequent gene expression was assessed (Fig. 6A). Samples either received a combination of the FC exposure and Nintedanib treatment, or FC exposure alone, which served as a control. Nintedanib treatment led to the upregulation of transitional epithelial markers *KRT17* (p = 0.0133) and *CLDN4* (p = 0.0145) relative to the FC-only samples (Fig. 6B). The alveolar epithelial type I marker *PDPN* was also upregulated (p = 0.0308), while the aberrant epithelial marker *FN1* [13, 63] was downregulated (p = 0.0196) in Nintedanib-treated samples (Fig. 6B). Meanwhile, expression levels of *SFTPC*, *CTGF*, *LAMP3*, and *AQP5* remained relatively unchanged between conditions (Fig. 6B).

**Fig. 6.**
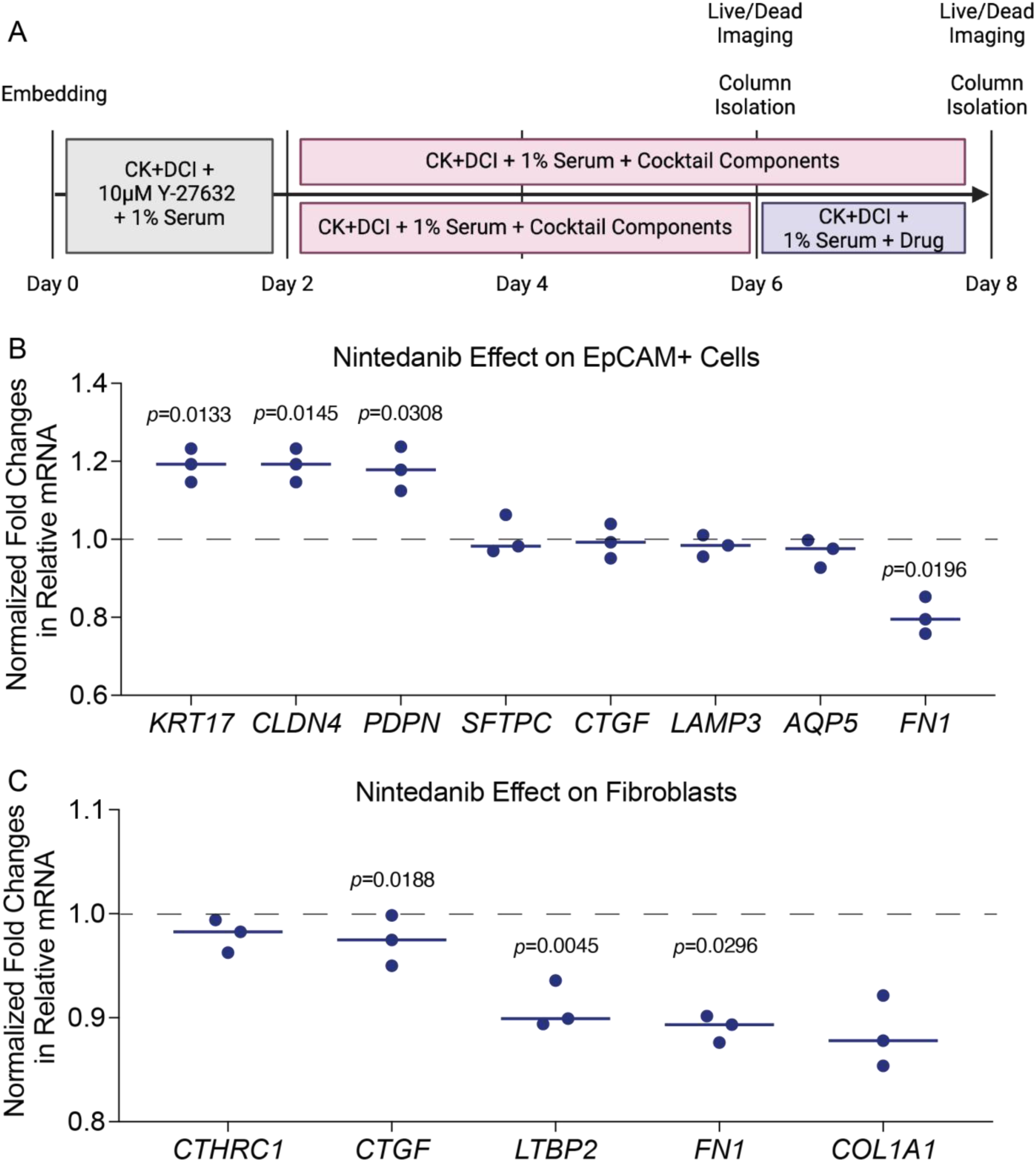
3D distal lung models were responsive to anti-fibrotic treatment. (A) Schematic representation of the therapeutic treatment experimental timeline and outputs. (B) Relative gene expressions for EpCAM^+^ cells after Nintedanib treatment, normalized to the FC only samples. Data presented as median with symbols representing biological replicates. Statistical significance was determined by paired t-tests between Nintedanib-treated and untreated conditions for each cell line. (C) Relative gene expressions for fibroblasts after Nintedanib treatment, normalized to the FC only samples. Data presented as median with symbols representing biological replicates. Statistical significance was determined by paired t-tests between Nintedanib-treated and untreated conditions for each cell line.

Similarly, several genes showed statistically significant differences when comparing Nintedanib-treated fibroblasts to FC-only fibroblasts. No fibroblast genes were upregulated in the Nintedanib-treated samples; however, *LTBP2* (p = 0.0188), *FN1* (p = 0.0045), and *COL1A1* (p = 0.0296) were all downregulated (Fig. 6C). Furthermore, expression levels of *CTHRC1 and CTGF* remained unchanged between conditions (Fig. 6C).

## Discussion

Here, we present novel 3D human lung models designed to support iATII- fibroblast co-culture within biomaterial systems that integrate synthetic, tunable stiffness hydrogels with patient-specific cells and serum as a model for pulmonary fibrosis [10].

By controlling the microenvironmental stiffness of the embedding hydrogel and delivering pro-fibrotic biochemical cues, we assessed how these factors synergistically drove epithelial injury and fibroblast activation. Given that nearly 70% of IPF cases occur in males [21], this study also aimed to establish a male-specific IPF model with male-derived iATIIs, fibroblasts, and serum to minimize sex as a confounding variable. Prior studies highlighted significant differences in fibroblast activation response based on the sex and age of the serum used to supplement the cell culture medium [64].

These results underscored the importance of using sex- and age-matched serum in disease modeling [64]. Expanding the availability of female iATIIs in the future and replicating this work with female cells and serum will be crucial for investigating potential sex-specific disease mechanisms and better understand the large dimorphism in this disease [65]. Likewise, the average age of onset for IPF is typically between 60 and 70 years old. It is rare in individuals under 50, and the risk increases with age [66]. The age of human serum also impacts relative hormone levels and thus cellular activation [64], so all human serum and fibroblasts used in this study were sourced from older (≥60 years old) patients to make the demographics of the populations most at-risk for IPF.

The hydrogel formulations presented in this study were designed to support iPSC-derived iATIIs, allowing for independent evaluation of microenvironmental stiffness and pro-inflammatory biochemical cues. Laminin, a key protein in the alveolar epithelium, plays a crucial role in lung morphogenesis and supports alveolar growth [67–69]. Laminin entrapped in hyaluronic acid hydrogels successfully supported ATII growth and self-assembly into spheroids without a need for Matrigel [70], but, to the best of our knowledge, this is the first study to replicate these results in a fully 3D engineered hydrogel system. One limitation of this work is that the laminin/entactin entrapped protein within the hydrogels was derived from mice. Identifying a suitable human laminin protein alternative would be beneficial and more translational to ensure the model consists entirely of human-derived materials. During lung development, MMP secretion predominantly shifts from MMP2 to MMP3 to MMP9 [46, 48], which reflects dynamic ECM remodeling and cellular behavior. MMP9 has also been widely implicated in early fibrotic tissue as critical ECM regulator, where it activates latent TGF-β1 and contributes to fibrosis progression [71–73]. Thus, the MMP9-degradable peptide crosslinker, laminin peptide mimic, and laminin/entactin protein complex were strategically included within the hydrogel formulations to facilitate cellular adhesion and remodeling in 3D [39].

Building on previously published work, a key distinction of this approach was that iATIIs naturally form spheres, or alveolospheres when cultured in 3D [34, 35, 70]. By leveraging this inherent geometry, iATIIs were aggregated successfully into a larger acinar structure without the need for microsphere templates [10, 44]. Photo-degradable microsphere templates have been used to generate cyst structures mimicking a single alveolus, a model which can be used to study crosstalk between templated epithelial cells and fibroblasts but which limits the ability of epithelial cells to provide autocrine support across a larger acinar structure, and also risks spontaneous and uncontrolled differentiation of primary cells [44, 45]. Self-assembling alveolospheres can also be generated in a 2.5D microwell system, where the dimensions of each well determine the ultimate spheroid size, but without full embedding in a supportive material [70]. Our 3D study allows for alveolospheres to be surrounded by tunable biomaterial and neighboring fibroblasts in a more physiologically relevant geometry. These engineered models also supported high total cell viability (>75%) throughout the 8-day culture period. While not statistically significant, a dip in cell viability to 78.17% ± 15.26% was observed on day 6 within the stiff hydrogels that were exposed to the fibrosis cocktail.

This trend suggests that the combined effects of increased microenvironmental stiffness and pro-fibrotic biochemical cues may have contributed to higher cell death. However, viability returned to above 80% by day 8, indicating that cell clearance may have occurred. This corresponds to observed effects of FC treatment on alveolospheres in matrigel, where exposure caused a trend towards increase in cellular damage markers without overall loss in number of alveolospheres [13], indicating that FC can cause a certain amount of stress without overt toxicity across a 3D culture.

Successful culture of iATII cells in engineered hydrogels is particularly advantageous given the widespread reliance of iATII growth in Matrigel [22, 31, 34, 35, 63, 70, 74], which presents significant regulatory challenges, as no Matrigel-derived products have been approved by the FDA or deemed safe for clinical applications such as autologous cell therapies [75]. Additionally, by incorporating a dynamic stiffening mechanism with the soft hydrogels, it would enhance the model’s physiological relevance and better recapitulate disease progression. Leveraging the tyrosine residues within the laminin, peptide mimics, and degradable crosslinker could enable ruthenium-based crosslinking, as demonstrated by Nizamoglu *et al.* [76–78]. Future work could also explore the use of hybrid-hydrogels, which have gained significant traction for harnessing the advantages of both synthetic and natural hydrogels [7, 14, 79].

Enhancing the hydrogel formulation with additional ECM proteins could improve viscoelasticity, increase cell remodeling, and further refine the model for studying fibrosis [77]. This design consideration will be particularly important if the goal is to seed single iATII cells within the hydrogels and allow them to self-assemble into alveolospheres over extended culture durations, such as several weeks.

In this study, the temporal expression of several ATII, transitional epithelial, ATI, and fibroblast activation genes were monitored to assess whether fibrotic markers could be effectively recapitulated. Treatment of alveolospheres alone in matrigel with FC has been observed to decrease expression of the ATII marker *SFTPC* while increasing the transitional marker *KRT17* and promoting aberrant ECM gene expression by epithelial cells, including expression of *FN1* [13]. Similarly, in our co-culture model, *SFTPC* [34, 35, 74] and *LAMP3* [23, 80] served as ATII-specific markers, while *KRT17* [23, 26, 29] and *CLDN4* [26, 81] identified transitional epithelial states. It is important to note that robust red fluorescence from the SFTPC^tdTomato^ reporter was observed at the time of seeding the iATIIs, indicating high activity of the *SFTPC* promoter [34, 35]. *PDPN* [13, 63, 74] and *AQP5* [22, 82] were used as ATI markers, while fibroblast activation was assessed through the expression of *COL1A1* [10, 83], *FN1* [84, 85], *CTGF* [86, 87], *CTHRC1* [83, 88], and *LTBP2* [89, 90]. Interestingly, CTHRC1^+^ cells have been shown to express pathologic ECM genes in fibrotic lungs and exhibit high mobility, often accumulating within fibrotic foci [83, 88, 91]. The upregulation of this gene within the FC-exposed samples suggests that early fibrosis may be occurring. Compared to the vehicle control samples, exposure to the fibrosis cocktail resulted in a clear and expected downregulation of ATII markers and upregulation in transitional epithelial, ATI, and fibroblast markers. However, *AQP5* exhibited a notable deviation from this trend, potentially suggesting that iATIIs may not have been able to fully differentiate into ATI cells or underwent apoptosis, leading to reduced cell survival.

The epithelial DOE also identified the hydrogel’s elastic modulus, the interaction between fibroblast donor age and elastic modulus, and fibrosis cocktail exposure as the top three factors to influence epithelial cell gene expression. In contrast, the fibroblast DOE ranked fibrosis cocktail exposure, the hydrogel’s elastic modulus, and time as the primary drivers of fibroblast activation. Notably, since all the input variables (age of the patient fibroblasts, elastic modulus, exposure, and time) were statistically significant within both DOEs, it provides strong evidence that these factors should be integrated into future fibrosis models.

Lastly, to further validate our model and assess its responsiveness to anti-fibrotic treatment, we tested the FDA approved drug Nintedanib. Alveolospheres in Matrigel treated with FC show an acquisition of mesenchymal-type markers including *FN1* that can be partially rescued by treatment with Nintendanib, but Nintendanib treatment in this model system fails to rescue *SFTPC* expression, indicating an incomplete reversion of epithelial cell injury [13]. In our co-culture system, after four days of treatment, *KRT17*, *CLDN4*, and *PDPN* were significantly upregulated relative to the FC-only control samples. This finding suggests that Nintedanib treatment in the context of fibroblast co-culture may have supported healthy ATII-to-ATI differentiation and preserved a AT1 cellular subpopulation. Additionally, *FN1* expression was evaluated in EpCAM^+^ cells, as this marker has been linked to the emergence of an aberrant basaloid cell population [13, 63]. Remarkably, Nintedanib downregulated *FN1* expression, which suggests that the epithelial cells were more likely to maintain normal functionality. In line with these favorable findings, Nintedanib treatment also downregulated *LTBP2*, *FN1*, and *COL1A1*, which are highly expressed in fibrotic tissue. These results were in line with a study that pretreated lung fibroblasts with Nintedanib before stimulation with

TGF-β and observed that Nintedanib protected fibroblasts from increased expression of fibrotic markers and phenotypes [92], though our timeline in which Nintedanib treatment follows fibrotic stimulation is more clinically relevant.

## Conclusion

In summary, we have developed an iATII-fibroblast model of human IPF that integrates key factors that contribute to fibrogenesis and evaluates respective impacts on gene expression. By leveraging tunable synthetic hydrogels, this biomaterial platform enables independent interrogation of biomechanical and biochemical cues. This model accurately reproduced the geometric and spatial cellular arrangement, as well as the mechanical properties of native distal lung tissue. While designed to simplify the complexity of fibrosis to the most critical parameters, our results demonstrate that donor age, hydrogel stiffness, pro-inflammatory biochemical cues, and time all significantly influence fibrotic gene expression. Our findings also identified that the combination of stiff hydrogels and fibrosis cocktail exposure resulted in the most fibrotic cellular microenvironment. Synergistically, these factors captured epithelial injury and fibroblast activation, which aligns these results with expected clinical outcomes. Furthermore, since Nintedanib treatment modulated fibrotic epithelial and fibroblast outcomes, it highlights the model’s potential for therapeutic screening.

Given the significant gap in our understanding of the events associated with fibrotic progression, we believe this model has the potential to address crucial mechanistic questions through systematic testing. The initial success of this model demonstrates feasibility that engineered hydrogels can support iATII growth in well-defined and controlled substrates that closely mimic the native lung environment. This work presents a more comprehensive *in vitro* distal lung model than what is currently available. By adopting an engineering approach, this work lays a solid foundation for further characterization of the cells at a protein level. Future studies should focus on developing female-specific human IPF models and incorporating dynamic stiffening and viscoelasticity into the hydrogels. Overall, this model serves as a powerful tool for studying IPF pathogenesis and evaluating anti-fibrotic interventions with strong physiological relevance.

## Data availability

The data that support the findings of this study are openly available in Mendeley Data at doi: 10.17632/3jchc2mkbf.1

## CRediT authorship contribution statement

**Alicia E. Tanneberger:** Conceptualization, Data curation, Formal analysis, Methodology, Project administration, Writing – original draft, Writing – reviewing & editing. **Rachel Blomberg**: Data curation, Formal analysis, Methodology, Writing – reviewing & editing. **Anton Kary**: Data curation, Methodology, Project administration, Writing – reviewing & editing. **Andrew Lu**: Data curation, Methodology, Writing – reviewing & editing. **David W.H. Riches**: Project administration, Supervision, Writing – reviewing & editing. **Chelsea M. Magin**: Conceptualization, Data curation, Formal analysis, Funding acquisition, Project administration, Supervision, Writing – original draft, Writing – reviewing & editing.

## Declaration of AI and AI-assisted technologies in the writing process

During the preparation of this work the author(s) used ChatGPT to improve readability and language. After using this tool/service, the author(s) reviewed and edited the content as needed and take(s) full responsibility for the content of the publication.

## Acknowledgements

The graphical abstract (Magin, C. (2025) https://BioRender.com/p27r262), portions of Fig. 1 (Magin, C. (2025) https://BioRender.com/t82x175), and part of Fig. 6 (Magin, C. (2025) https://BioRender.com/p47v260) were created using BioRender. The authors sincerely thank Dr. Darrell Kotton (Boston University) and the Center for Regenerative Medicine at Boston University for providing iATIIs and technical support that were instrumental to these experiments. We also acknowledge Dr. David W.H. Riches (National Jewish Health) and Benjamin Edelman (National Jewish Health) for their assistance in coordinating and acquiring patient-specific fibroblasts. Additionally, we are grateful to Dr. Amy L. Ryan (University of Iowa) for her valuable technical input and feedback that guided these experiments in its early stages. We further acknowledge Mikala M. Mueller (Magin Lab, CU Denver | Anschutz) for her support with medium changes and chemistry, as well as Dema Essmaeil (Magin Lab, CU Denver | Anschutz) for her early work in optimizing iATII-fibroblast aggregations.

