## Supplementary Material for "Biomaterial-based 3D human lung models replicate pathological characteristics of early pulmonary fibrosis"

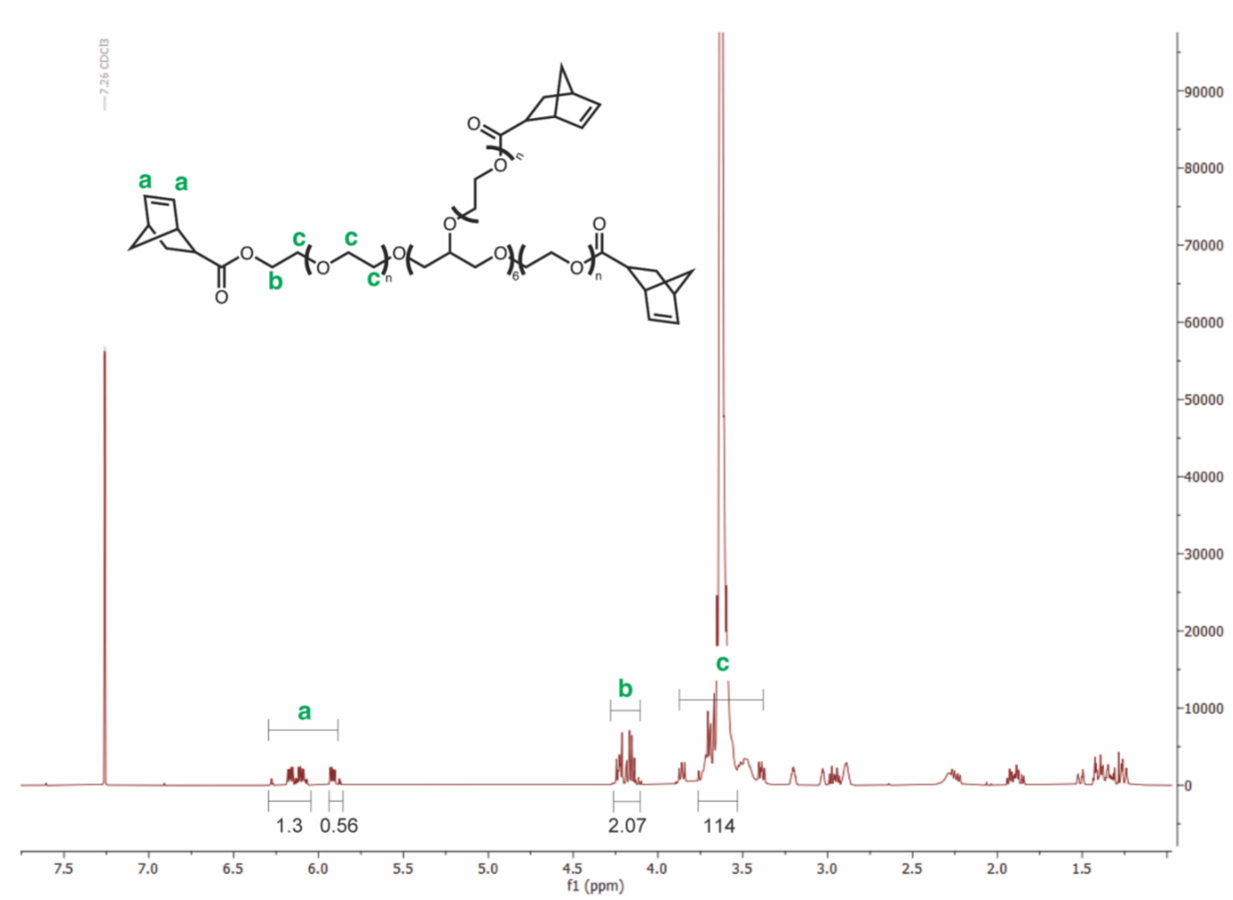


**Fig. S1.** $^1$H NMR results of 10 kg/mol PEGNB (Bruker DPX-400 FT NMR spectrometer). δH (ppm) (300 MHz, $CDCl_3$): 3.71 (s, 114H, PEG $CH_2$-$CH_2$), 4.1-4.2 (m, 2H, -$CH_2$-O), 5.9-6.2 (m, 2H, –CH=CH-). NMR shows a 93% quantitative norbornene functionalization based on a comparison of the alkene protons from norbornene to the theoretically expected number of alkene protons, using the ethylene glycol protons as a reference.

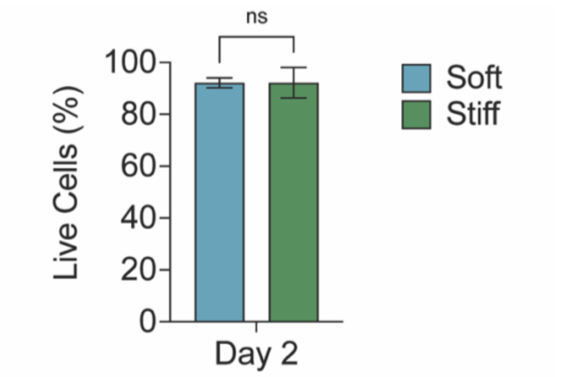


**Fig. S2.** iATIIs and fibroblasts maintained high cell viability during the first 48 hours after being embedded in hydrogels. The percentage of live cells was quantified using a

ReadyProbes Cell Viability Imaging Kit. Statistical significance was determined by a paired t-test (n=6, ns = no significance).

**Table S1.** CK-DCI media composition

| Component | Final amount |
|---|---|
| IMDM | Base Media-75% v/v |
| Ham's F-12 | Base Media-25% v/v |
| B27 (50x) | 0.5x |
| N2 (100x) | 0.5x |
| GlutaMax (100x) | 1x |
| BSA Fraction V | 0.05% |
| Ascorbic acid | 50 µg/ml |
| 1-Thioglycerol | 450 µM |
| Primocin | 1x |
| CHIR99021 | 3 µM |
| KGF | 10 ng/µl |
| Dexmethasone | 50nM |
| cAMP | 0.1mM |
| IBMX | 0.1mM |

**Table S2.** Human primers for RT-qPCR

| Gene | Forward 5’-3’ Primer Sequence | Reverse 5’-3’ Primer Sequence |
|---|---|---|
| RPS18 | ATGCAGAATCCACGCCAGTA | TTCCATCCTTTACATCCTTCTGTCT |
| SFTPC | GCTCCAGAGAGCATCCCCAG | GCCTGCAGAGAGCATTCCAT |
| LAMP3 | CGGGCATTCCTTCAAGTGCG | AGGCAGAGACCAACCACGAT |
| KRT17 | CAGTCCCAGCTCAGCATGAA | CCACAATGGTACGCACCTGA |
| CLDN4 | TCTGCTCACACTTGCTGGCT | CATTGTTCAGCGTCCACGGG |
| PDPN | ACAGGCATTCGCATCGAGGA | GGCGAGTACCTTCCCGACAT |
| AQP5 | ACTGGCTGCTCCATGAACCC | TGATGGCCACACGCTCACT |
| RPL30 | TCTGCTTGTACCCCAGGACG | CGAAAAAGTCGCTGGAGTCGAT |
| COL1A1 | TCTGCGACAACGGCAAGGTG | GACGCCGGTGGTTTCTTGGT |
| FN1 | AGGAAGCCGAGGTTTTAACTG | AGGACGCTCATAAGTGTCACC |
| CTGF | AGGAGTGGGTGTGTGACGA | CCAGGCAGTTGGCTCTAATC |
| CTHRC1 | CAGACGCTGACCACGTTCC | GCATTTTAGCCGAAGTGAGCC |
| LTBP2 | GGATGGACAACAGCAAACAGC | CATCGGGAATGACCTCCTCG |
